# A novel mouse model of hypertensive emergency with multiorgan microvascular disease implicating the VEGFA/sFlt-1 balance

**DOI:** 10.64898/2026.03.03.709451

**Authors:** Thibaut d’Izarny-Gargas, Imane Bensaada, Christophe Roubeix, Léa Guyonnet, Véronique Baudrie, Sarah Azancot, Pauline Maurissens, Léa Resmini, Arthur Lavigne, Christine Ibrahim, Léa Dionet, Anna Chipont, Carole Hénique, Philippe Bonnin, Xavier Guillonneau, Jérôme Thireau, Florian Sennlaub, Neeraj Dhaun, Olivia Lenoir, Pierre-Louis Tharaux

**Affiliations:** Université Paris Cité, Inserm, PARCC, F75015 Paris, France; Cytometry platform, Curie Coretech, Institut Curie; Sorbonne University, INSERM, CNRS, Institut de la Vision, 17 rue Moreau, F-75012 Paris, France; Centre Hospitalier National d’Ophtalmologie des Quinze-Vingts, INSERM-DHOS CIC 503, Paris, France; Hôpital Européen Georges Pompidou, Pharmacologie Clinique, Assistance Publique Hôpitaux de Paris, Paris, France; Université Paris-Est Créteil, Créteil, F-94010 France; Institut National de La Santé Et de La Recherche Médicale (INSERM) U955, Institut Mondor de Recherche Biomédicale (IMRB), Créteil, F-94010 France; Physiologie clinique - Explorations fonctionnelles, Hôpital Lariboisière, APHP, Paris, France; Université de Montpellier, CNRS UMR 9214, INSERM U1046, PhyMedExp, Montpellier, France; Edinburgh Kidney, Institute for Neuroscience and Cardiovascular Research, University of Edinburgh, Edinburgh, UK; Department of Renal Medicine, Royal Infirmary of Edinburgh, Edinburgh, UK

## Abstract

**Background:** Hypertensive emergency (HTEM) is defined by abrupt blood pressure elevation with acute multi-organ damage, yet the mechanisms predisposing only a subset of hypertensive individuals to HTEM remain unclear. Progress has been limited by the lack of a mouse model that faithfully replicates human disease. We aimed to identify determinants of susceptibility to hypertensive microvascular injury and characterize a murine model of HTEM.

**Methods:** Male C57BL/6J (B6J) and 129S2/SvPasCrl (129Sv) mice were exposed to severe hypertension via angiotensin II infusion combined with a high-salt diet. We assessed survival, renal and retinal injury, cardiac function and electrophysiology, vascular permeability, circulating angiogenic factors, and glomerular transcriptional profiles using single-cell RNA sequencing. Bone marrow transplantation and recombinant human PlGF-2 treatment were used to investigate mechanisms driving endothelial injury.

**Results:** Despite comparable blood pressure, 129Sv mice, but not B6J, developed malignant hypertension with albuminuria, acute kidney injury, retinal hemorrhages, microvascular leakage, cardiac dysfunction, and arrhythmias. Hypertensive 129Sv mice exhibited markedly elevated circulating sFlt-1. PlGF-2 supplementation partially reversed albuminuria, preserved glomerular ultrastructure, and reduced retinal hemorrhages. Bone marrow transfers revealed contributions from both hematopoietic and non-hematopoietic 129Sv compartments to sFlt-1 overproduction and organ injury. Single-cell transcriptomics revealed profound repression of angiogenic, metabolic, and stress-response pathways in glomerular endothelial cells, a repression partially restored by PlGF-2.

**Conclusions:** We identify 129Sv mice as a robust model of HTEM, exhibiting multi-organ microvascular injury that closely mirrors the human condition. Our results reveal blood-pressure-independent susceptibility to organ damage and implicate dysregulated VEGFA/sFlt-1 signaling as a central driver of endothelial dysfunction, highlighting angiogenic imbalance as a potential therapeutic target.

## Introduction

Hypertension is recognized as the most prevalent modifiable risk factor worldwide for both mortality and cardiovascular morbidity (1). Chronically elevated blood pressure (BP) is associated with progressive impairment in target organs, including the heart, brain, and kidneys, and long-term treatment of hypertension has been demonstrated to lower the incidence of mortality, major adverse cardiovascular events, and dementia, and to slow the progression of chronic kidney disease (2). A fraction of hypertensive patients, however, may present with severely elevated BP and acute development of Hypertension Mediated Organ Damage (HMOD) or target organ damage (TOD), defining hypertensive emergency (HTEM), a syndrome previously known as malignant hypertension (3–6).

HTEM is a systemic disease featuring multi-organ damage occurring acutely in the context of severe hypertension (7). Various diagnostic criteria have been proposed for this condition. Rather than a specific BP threshold, modern definitions of HTEM insist on the presence of high- grade retinopathy and/or TOD as the primary diagnostic features (4, 8). While the short-term prognosis of HTEM has been improved with modern antihypertensive medication, long-term follow-up demonstrated that these patients remain at a high risk of cardiovascular disease and end-stage kidney disease (9, 10). Improving the long-term prognosis of HTEM patients may require a deeper understanding of this condition’s pathophysiology. At similar BP levels, most patients with severe hypertension will not develop acute TOD, while only a small proportion will present with HTEM. This suggests that BP-independent mechanisms play a major role in the disease development. Likewise, in HTEM patients, prognosis is more closely tied to organ impairment than to initial BP level (11, 12).

Rat models of malignant hypertension, such as the stroke-prone spontaneously hypertensive rat and the transgenic *Ren2* Sprague-Dawley rat, have highlighted the central role of the renin-angiotensin-aldosterone system in the pathophysiology of kidney lesions (13, 14). This is illustrated in the Kincaid-Smith model of malignant hypertension, which proposes that severe hypertension and endothelial damage result in kidney ischemia and a subsequent vicious cycle of renin-dependent BP elevation (15). However, the extent to which renin elevation accounts for HTEM in human patients remains unclear, as a significant proportion of these patients present with normal or comparable plasma renin activity to patients with chronic, non-malignant hypertension (16, 17). Furthermore, primary hyperaldosteronism with suppressed plasma renin activity can be complicated by HTEM (18, 19). Moreover, hypertensive patients with a past episode of HTEM and controlled BP exhibit persistent vascular dysfunction (20), suggesting that additional, vessel-centric factors should be sought out to better explain the occurrence of acute TOD after BP elevation.

To the best of our knowledge, no mouse model of hypertension that recapitulates the features of human HTEM is currently available. The commonly used C57BL/6J (“B6J”) mouse is considered resistant to hypertensive kidney injury; however, other strains may be more susceptible to kidney damage (21). In this work, we show that 129S2/SvPasCrl (“129Sv”) mice challenged with severe hypertension exhibit a lethal phenotype that is highly reminiscent of human HTEM. We further demonstrate the mechanistic role of an alteration in angiogenesis balance in this model, which may open new avenues for physiopathological and therapeutic investigations in human patients.

## Methods

### Animals

Wild-type C57BL6/J (B6J) and 129S2/SvPasCrl (129sv) were obtained from Charles River Laboratories. Experiments were conducted in accordance with the French veterinary guidelines and with the European Community guidelines for the use of experimental animal (L358–86/609EEC). They were approved by the Institut National de la Santé et de la Recherche Médicale and the local University Research Ethics Committee (CEEA34 Paris Cité University, Apafis #7646 and #19592). Mice underwent sanitary control tests to confirm pathogen-free status and were housed in the same animal facility before any experiments. All animal procedures were conducted in accordance with ARRIVE guidelines. Animals were housed under standard laboratory conditions with controlled temperature and humidity, and a 12- hour light/dark cycle. Environmental enrichment was provided. Endpoints were established, and animals were monitored daily for signs of distress. The experimental unit was a single animal. Sample sizes were determined based on prior studies and pilot data; no formal power calculation was performed. All animals were randomly assigned to treatment or control groups. Investigators were blinded to group allocation during data analysis. No animals or data points were excluded from analysis. The number of animals used is indicated in each figure legend.

### Hypertension model

The hypertensive model was induced by subcutaneous infusion of angiotensin II (Ang II) (Alomone, GPA-100) at 1 µg/kg/min for 7 or 14 days, via osmotic minipumps (Alzet, model 1007D or 1002) in 10- to 16-week-old males. Mice were anesthetized with isoflurane and pumps were implanted subcutaneously on the back between the shoulder blades and hips. Subcutaneous dose of buprenorphine (0.05mg/kg) was administered during the surgery and twice a day for 72 hours. Mice received high-salt diet (3% NaCl in food, SSNIFF Spezialdiäten GmbH) and were euthanized 7 or 14 days after implantation of minipumps. Recombinant human placental growth factor (PlGF)-2 PlGF-2 (rhPlGF-2) (PeproTech) was delivered to hypertensive 129Sv mice at a dosage of 80 µg/kg/day using a 14-day osmotic minipump (Alzet, model 1002) which was implanted subcutaneously during the same procedure as Ang II minipump implantation.

### Telemetry blood pressure (BP) measurement

Mice were anesthetized with isoflurane and a BP telemeter (model TA11PA-C10, Data Sciences International) was implanted in the left femoral artery. A single dose of amoxicillin (20 mg/kg administered intraperitoneally, Clamoxyl ; SmithKlineBeecham) and ketoprofen (5 mg/kg administered intraperitoneally, Profenid ; Aventis) was administered after the surgery. BP was measured in conscious, unrestrained mice. Recordings were obtained every 15 minutes for 60 seconds; daytime was defined as 07:00–19:00 and nighttime as 19:00–07:00. Mice were adapted for 1 week, and BP values from the last 3 days of the adaptation period were averaged to define the basal BP.

*Bone marrow transplantation*.

We conducted bone marrow transplantation experiments, leveraging the congenic markers cluster of differentiation (CD) 45.1 and CD45.2. 8-week-old male 129Sv and B6J CD45.2 mice were irradiated (2 x 6.5G, 4 hours apart) and reconstituted with an intravenous (i.v.) injection in the retro-orbital vein of 10^7^ bone marrow cells isolated from femurs of either 129Sv or B6J CD45.1 mice. Mice with correct engraftment were considered when donor cells accounted for ≥85% of CD45⁺ leukocytes at 6 weeks. After 6 weeks of recovery, these mice were infused with Ang II.

### Assessment of renal function and albuminuria

All mice were placed in metabolic cages with free access to water for a 6-hour urine collection. Urinary creatinine, plasma albumin, and plasma urea concentrations were spectrophotometrically analyzed by using a colorimetric method (Olympus, AU400). Urinary albumin excretion was measured using a specific enzyme-linked immunosorbent assay for the quantitative determination of albumin in mouse urine (Crystal Chem, 80630).

### Renal histopathology

For optical microscopy, kidneys were immersed in 10% formalin and embedded in paraffin. Four-millimeter sections were processed for histopathology and stained with Masson’s trichrome. Photomicrographs were taken using a Zeiss Axiophot photomicroscope (Jena, Germany). For electron microscopy, sections of renal cortex were fixed in Trump’s solution (EMS, 11750) and embedded in Araldite M (Sigma Aldrich, 10951). Ultrathin sections were counter-stained with uranyl acetate and lead citrate and examined with a transmission electron microscope (JEM-1011, JEOL, Peabody, MA).

### Assessment of retinal injury

In Life imaging: Before anesthesia, pupils were dilated with tropicamide (Mydriaticum®, Théa, France) and phenylephrine (Néosynephrine®, Europhta, France). Animals were then anesthetized by isoflurane inhalation (Axience, France) and placed first in front of the Micron 3 device (Phoenix Research Labs, Oregon, USA) for eye fundus imaging. Then, animals were transferred in front of the SD-OCT imaging device (Bioptigen, 840nm HHP; Bioptigen, North Carolina, USA) where anesthesia was maintained with a mask for continuous isofurane inhalation. Eyes were kept moist with 0.9 NaCl throughout the entire procedure. Image artifacts due to respiration were first eliminated using the StackReg Plugin. Then, each movie was converted into a single image by compiling a Z-projection of all the images. Eye fundus and OCT imaging were performed in C57BL/6J and 129/Sv mice prior to and following 7 days of Ang II. Retinal detachment amplitude was quantified for each eye using an OCT-based scoring system: Grade 0 (no detachment); Grade 1 (< 25 µm); Grade 2 (26–50 µm); Grade 3 (51–75 µm); Grade 4 (76–100 µm); and Grade 5 (> 100 µm).

After enucleation, the eyes were fixed in 4% paraformaldehyde for 30 minutes at room temperature. After washing in phosphate-buffered saline and removal of the cornea and lens, the retina was dissected from the retinal pigment epithelium/choroid/sclera, and haemorrhagic spots were counted. Retinas then were incubated overnight at 4C antibodies with primary (lectin Alexa conjugated, 1:100, Invitrogen) in phosphate-buffered saline supplemented with 0.5% Triton X-100). Retinas were flat-mounted and viewed with a fluorescence microscope (DM5500B, Leica, France). Vascular networks were calculated from an average of 8 images/animal (4/retina). Vascular network density was obtained using the NeuronJ plugin on ImageJ software.

*Noninvasive ultrasound assessment of cardiac and renal hemodynamics* Ultrasound examination was performed under isoflurane anesthesia with 0.75% Isoflurane in 100% O2, during spontaneous breathing, using an echocardiograph (Acuson S3000 ; Siemens, Erlangen, Germany) equipped with a 14 MHz linear transducer (14L5 SP). Mice were placed on a heating blanket (38 °C) to avoid hypothermia. A parasternal long-axis B-mode image was used to measure cardiac output; the animal was placed in left lateral decubitus, and the ultrasound device was placed on the anterior part of the animal’s chest for cardiac output acquisition. These positions avoid placing pressure on the abdomen or on the chest with the transducer and allow the visualisation of the pulmonary artery with the possibility to measure the inner diameter and the time-averaged mean blood flow velocity for cardiac output calculation. Cardiac output was calculated from the mean of 3 successive measurements. Two-dimensional parasternal long-axis views of the left ventricle (LV) were obtained for guided M-mode measurements at the end of the diastole (d) and systole (s). For functional analysis of the LV viable myocardium, we evaluated the LV diastolic and systolic internal diameter (LVIDd, LVIDs). Percentage fractional shortening (%FS) was calculated by the following formula: %FS= [(LVIDd-LVIDs)/LVDd]x 100. Heart dimensions measured, in both systole and diastole, included left ventricular posterior wall thickness, left ventricular internal diameter, and interventricular septum thickness. Stroke volume was calculated by dividing the CO by the HR.

The ultrasound device was then placed on the left side of the abdomen of the animal for ultrasound analysis of the kidneys and the right renal artery. A pulsed Doppler sample gate was placed on the longitudinal axis of the right renal artery, and the pulsed Doppler spectrum was recorded. Time-averaged mean blood flow velocity was calculated with correction of the angle between the long axis of each vessel and the Doppler beam.

*Electrocardiograms:* ECGs were recorded and analyzed off-line using the Ponemah Physiology Platform (Data Sciences International, USA) (22) in accordance with the Lambeth conventions for determining arrhythmic events (23). Under anesthesia (2.5% isoflurane/O_2_, Aerane), mice were implanted with subcutaneous devices (Physiothel TAE-F10 model, Data Sciences International, USA) and ECG signals recorded on days 0, days 2, and days 12 (1000Hz-sampling rate). Heart rate (bpm), corrected QT (QTc) intervals, and ST elevation (STE) were calculated using 3h ECGs (>60,000 identified PQRS complexes), taking in the same time slots at days 0 (baseline), 2, and 12. SDNN was calculated as a global variability index. The QT interval was defined from lead II of the ECG as the time between the first deviation from an isoelectric PR interval to the return of the ventricular repolarization to the isoelectric baseline. This method accommodates the low-amplitude portion of the T-wave and allows for complete ventricular repolarization (22). The end of the T-wave was automatically assessed by software and validated by hand by the same operator throughout the experiment. The QT correction was performed using the adapted Bazett’s formula by Mitchell (24). The ST elevation, a hallmark of myocardial suffering, was automatically assessed by the Ponemah Physiology Platform. The triggering of sinus arrest, atrioventricular block, atrial fibrillation or flutter, ventricular extrasystoles and tachycardia, sustained ventricular tachycardia, as defined by the Lambeth conventions, were identified and counted.

### Assessment of vascular permeability

Blood was sampled through intracardiac puncture with EDTA-coated syringes at the time of euthanasia, and a complete blood cell (CBC) count was performed with a Hemavet 950 cell counter 10 (Drew Scientific). Hearts and kidneys were harvested, weighed to determine wet weight, dried in a 60°C convection oven for 48 hours, and then weighed again to determine dry weight. Wet-to-dry weight ratios were computed.

### Evans Blue Permeability Assay

Evans Blue dye (Sigma-Aldrich Inc., St Louis, MO, USA) (0.5%) was administered intravenously via retro-orbital injection at a dose of 4 µL/g of mouse body weight under isoflurane (3%) anaesthesia. Two hours post-injection, mice were euthanized, the right atrium was dissected, and PBS was perfused through the apex of the left ventricle to flush out blood from the vasculature. Organs, including the heart, kidneys, one lung lobe, one liver lobe, and the spleen, were harvested and weighed. Each organ was incubated in formamide at 55°C for at least 48 hours. The volume of formamide used was adjusted according to the organ’s weight. During incubation, Evans Blue dye diffused from the tissue into the formamide. To quantify dye concentration, formamide absorbance was measured at 620 nm and 740 nm using the SoftMax software on a FlexStation 3 (Molecular Devices). Vascular permeability was expressed as absorbance normalized to organ weight.

### sFLT1 ELISA

Plasma soluble fms-like tyrosine kinase-1 (sFlt-1) was measured using enzyme-linked immunosorbent assay (MVR100, Biotechne) according to the manufacturer’s instructions. Samples were analysed in duplicate at 450nm with wavelength correction at 570nm (Synergy HT BioTEK). Concentrations were calculated from a standard curve using second-order (quadratic) regression analysis.

### Flow cytometry

Bone marrows were flushed out from leg bones and filtered using 40 µm cell strainers. Bone marrow suspension was treated with red blood cell lysis buffer (Sigma, #R7757) for 10 min at room temperature. Cells were counted, resuspended in staining buffer (PBS, 10% BSA and 2mM EDTA) and stained using anti-mouse CD45.1 (clone A20, BD), anti-mouse CD45.2 (clone 30-F11, BD), and viability dye for 30min in the dark at 4°C. Cells were centrifuged at 500g for 5 min, washed twice with staining buffer, and resuspended in staining buffer for flow cytometry analysis. Cells were acquired on an LSRII instrument (Becton Dickinson). Supervised analysis was performed using FlowJo software v10 (FlowJo LLC).

### Glomerular preparation for single-cell RNA sequencing

Mouse kidneys were minced and digested with collagenase IV (2 mg/ml) in calcium- and magnesium-free HBSS at 37 °C for 3 minutes. Digestion was stopped by adding cold HBSS supplemented with 10% FCS. The tissue suspension was filtered through a 100 µm mesh, and residual fragments were gently pressed through using a syringe plunger. After rinsing with cold HBSS, the suspension was passed through a 40 µm mesh that retained glomeruli while allowing smaller tubular fragments to pass through, thereby enriching the sample for glomeruli. Glomeruli were further purified by differential adhesion on a culture dish, allowing tubular fragments to settle (25). The enriched glomeruli were centrifuged at 290 g for 5 minutes at 4 °C. The pellet was resuspended in enzyme solution (2 mg/ml collagenase IV + 20 U/ml DNase in HBSS) and incubated at 37°C for 20 minutes with gentle trituration every 5 minutes. Digestion was stopped with HBSS containing 10% FCS, and the suspension was filtered through a 40 µm mesh. Cells were centrifuged at 400 g for 4 minutes at 4 °C and washed twice in HBSS supplemented with 1% FCS and 1 mM EDTA. Cell concentration was determined, and cells were loaded onto the 10x Genomics Chromium instrument.

### Single-cell library generation, sequencing, and data integration

Single-cell Gel Bead-In-EMulsions (GEMs) were generated using a Chromium Controller instrument (10x Genomics). Sequencing libraries were prepared using Chromium Single Cell 3’ Reagent Kits v3.1 (10x Genomics), according to the manufacturer’s instructions. Briefly, GEM-RT was performed in a thermal cycler at 53°C for 45 min, followed by 85°C for 5 min. Post GEM-RT Cleanup using DynaBeads MyOne Silane Beads was followed by cDNA amplification (98°C for 3 min, cycled 11 x 98°C for 15 s, 67°C for 20s, 72°C for1 min, and 72°C 1 min). After a cleanup with SPRIselect Reagent Kit and fragment size estimation with High Sensitivity™ HS DNA kit runned on TapeStation Bioanalyzer (Agilent), the libraries were constructed by performing the following steps: fragmentation, end-repair, A-tailing, SPRIselect cleanup, adaptor ligation, SPRIselect cleanup, sample index PCR, and SPRIselect size selection. The fragment size estimation of the resulting libraries were assessed with High Sensitivity™ HS DNA kit runned on TapeStation Bioanalyzer (Agilent) and quantified using the Qubit™ dsDNA High Sensitivity HS assay (ThermoFisher Scientific). Libraries were then sequenced in pairs on a HighOutput flow cell using an Illumina Nextseq 2000 with the following mode: 28 bp (10X Index + UMI), 10 bp (i7 Index), 10 bp (i5 Index), and 90 bp (Read 2). Fastq files were then uploaded to the Cell Ranger Count pipeline for demultiplexing, alignment, counting, and clustering (CellRanger: version-9.0.1, Mouse reference transcriptome: GRCm39-2024-A). The integrated dataset was exported and imported into R as a Seurat v5 object for downstream analysis (26). The integrated Seurat object was loaded using the qs package. Cells were annotated based on sample identifiers and grouped into three experimental conditions: control (129), Ang II-treated (129_Ang2), and Ang II + rhPlGF-2 treated (129_Ang2_PlGF). To optimize memory usage, Seurat layers were joined before subsetting. The RNA assay was used for normalization (NormalizeData), identification of highly variable genes (FindVariableFeatures), scaling (ScaleData), and dimensionality reduction via PCA (RunPCA). Clustering was performed using FindNeighbors and FindClusters (resolution = 0.5), followed by UMAP embedding (RunUMAP). Cells were retained if they expressed between 200 and 6000 genes and had mitochondrial gene content below 80%. Genes expressed in fewer than 10 cells were excluded from downstream analyses.

Cell types were annotated using module scores derived from canonical kidney markers for podocytes (*Nphs1*, *Nphs2*, *MaP*, *Wt1*, and *Podxl*), mesangial cells (*Pdgfrb*, *Pdgfa*, *Acta2*, *Itga8*, *Gata3*, and *Rgs5*), and endothelial cells (*Kdr*, *Pecam1*, *Cdh5*, *Emcn*, and *Klf2*). Cells were assigned to a major cell type if their module score exceeded the 90th percentile threshold. Doublets were identified by detecting co-expression of mutually exclusive markers across cell types and removed from the dataset.

Pairwise differential expression (DE) analyses were performed between experimental groups within each major cell type using Seurat’s FindMarkers function (Wilcoxon rank-sum test). Genes were considered significantly differentially expressed if adjusted p-values were < 0.05 and absolute log2 fold change > 0.5. Significantly up- or downregulated genes were subjected to pathway enrichment analysis using Gene Ontology (GO), HALLMARK gene sets (MSigDB) (27), and Reactome pathways (28). Enrichment was performed using enrichGO, enricher, and enrichPathway functions from the clusterProfiler (29), msigdbr, and ReactomePA packages.

### Statistical analyses

All graphs represent individual values and mean ± SEM. Statistical analyses were performed using GraphPad Prism software, version 9. Comparison between 2 groups was performed using a parametric Student t test when samples passed the Anderson-Darling and D’Agostino normality tests and the F test for equality of variance. Otherwise, a nonparametric Mann-Whitney test was used. Values of P < 0.05 were considered significant. ∗P < 0.05, ∗∗P < 0.01, ∗∗∗P < 0.001. Comparisons between multiple groups were performed using 1-way or 2-way ANOVA or Welch ANOVA when the SDs were significantly different between groups. Post hoc test was applied using Tukey or Dunnett’s T3 correction.

## Results

### Ang II + high-salt diet-induced hypertension induces mortality in 129Sv mice

First, we assessed mortality in wild-type B6J and 129Sv males under hypertensive conditions. While no death was recorded in B6J mice, lower survival was observed in 129Sv mice starting about a week after hypertension onset (Fig. 1A). Telemetry BP measurement showed similar values between mouse strains (Fig. 1B-C), demonstrating the observed mortality difference is due to poorer tolerance to elevated BP in 129Sv males, rather than heightened sensitivity to Ang II hypertensive effect.

**Figure 1:**
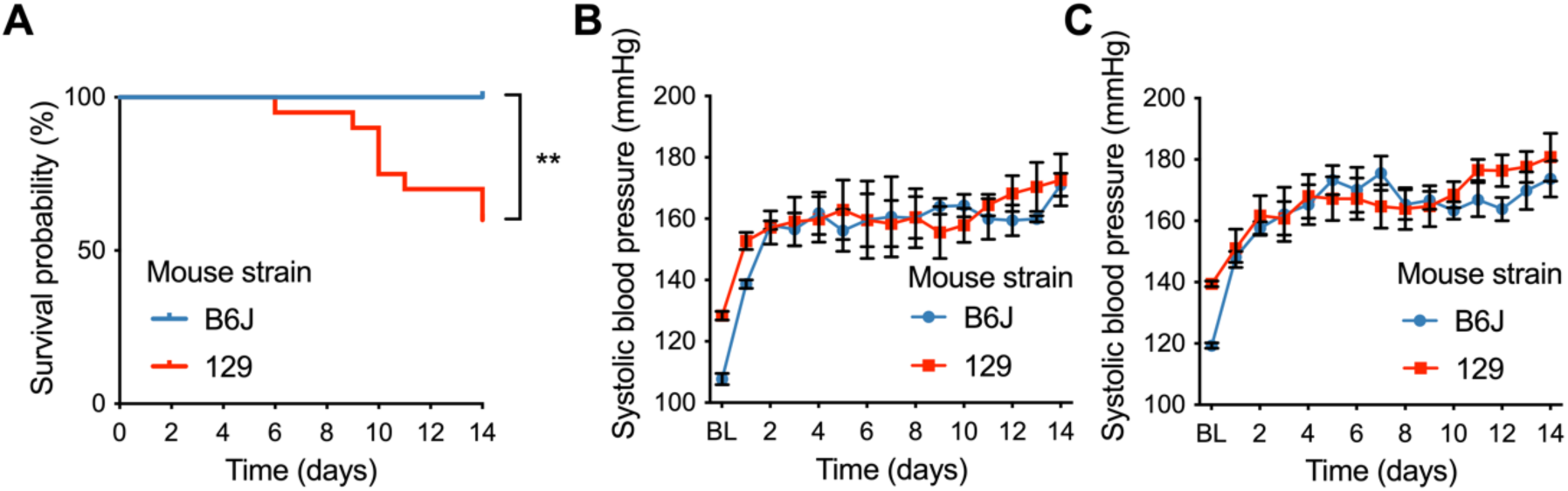
Angiotensin-2-induced hypertension promotes mouse death in the 129Sv strain but not in the C57BL6/J strain. A- Mice survival after hypertension onset (days after beginning of Ang II infusion). N= 20 C57BL6/J (B6J) and 129Sv (129) B- Mean daytime telemetry blood pressure N= 4 B6J and 129Sv. Values represent mean ± SEM. C- Mean nighttime telemetry blood pressure. N= 4 B6J and 129Sv. Values represent mean ± SEM.

### 129Sv hypertensive mice exhibit kidney and retinal injury suggestive of systemic microvascular disease

High blood pressure markedly increased urinary albumin excretion (ACR) in hypertensive 129Sv mice with progressive worsening over the 14-day timespan, while B6J mice remained free of significant albuminuria (Fig 2A). At day 7 after hypertension onset, we observed lower plasma albumin in 129Sv mice (Fig 2B), consistent with urinary albumin loss, and elevated blood urea nitrogen (BUN) (Fig 2C), consistent with acute kidney injury. Accordingly, histopathological examination of hypertensive 129Sv mice revealed tubular and glomerular filtration barrier (GFB) injury, including podocyte foot process effacement and endotheliosis (Fig 2D).

**Figure 2:**
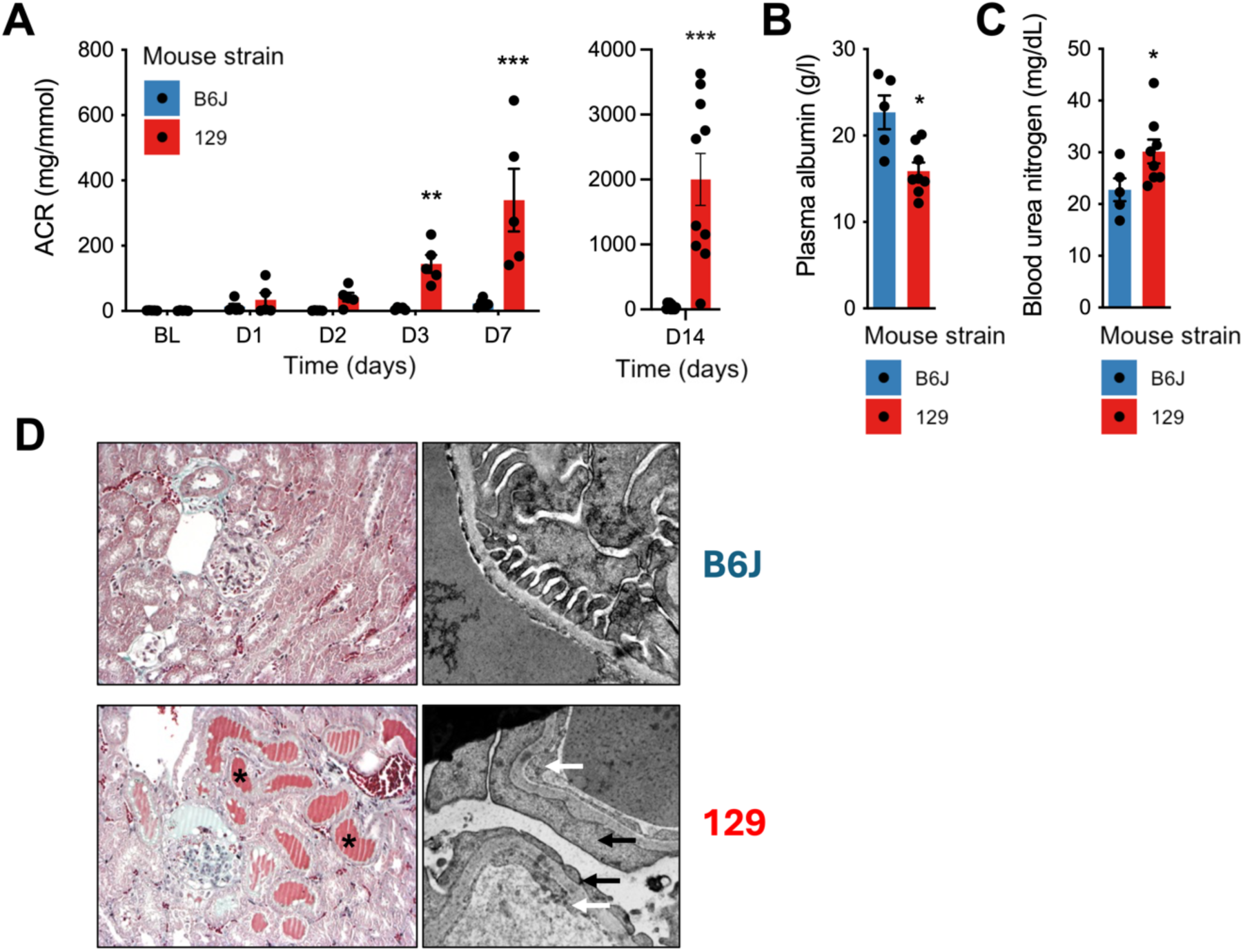
The 129Sv strain develops malignant hypertension with renal injury. A- Urine albumin-to-creatinine ratio across a 14-day hypertensive challenge. N=5 mice per group from day (D) 0 to D7 and N= 10 for D14. Values represent mean ± SEM. ** p < 0.01. B- Plasma albumin measured at day 7 of hypertensive challenge. N=5 C57BL6/J and 8 129Sv mice. Values represent mean ± SEM. *** p < 0.001. C- Blood urea nitrogen measured at day 7 of hypertensive challenge. N=5 C57BL6/J and 8 129Sv (129) mice. Values represent mean ± SEM. * p < 0.05. D- Representative photomicrographs of kidney histology on Masson’s trichrome staining (left panels) and glomerular filtration barrier ultrastructure in electronic transmission microscopy (right panels) at day 7 of hypertensive challenge, demonstrating tubular injury (asterisks), endothelial swelling and loss of fenestrations (white arrows), and podocyte foot process effacement (black arrows).

Because albuminuria and endotheliosis were strongly suggestive of microvascular impairment, we assessed retinal phenotype in mice at day 7 after the onset of hypertension. Consistent with our hypothesis, hypertensive 129Sv mice displayed retinal hemorrhagic spots (Fig. 3A-B), subretinal swelling (Fig. 3C) as well as capillary rarefaction (Fig. 3D and Supp. Fig. 1).

**Figure 3:**
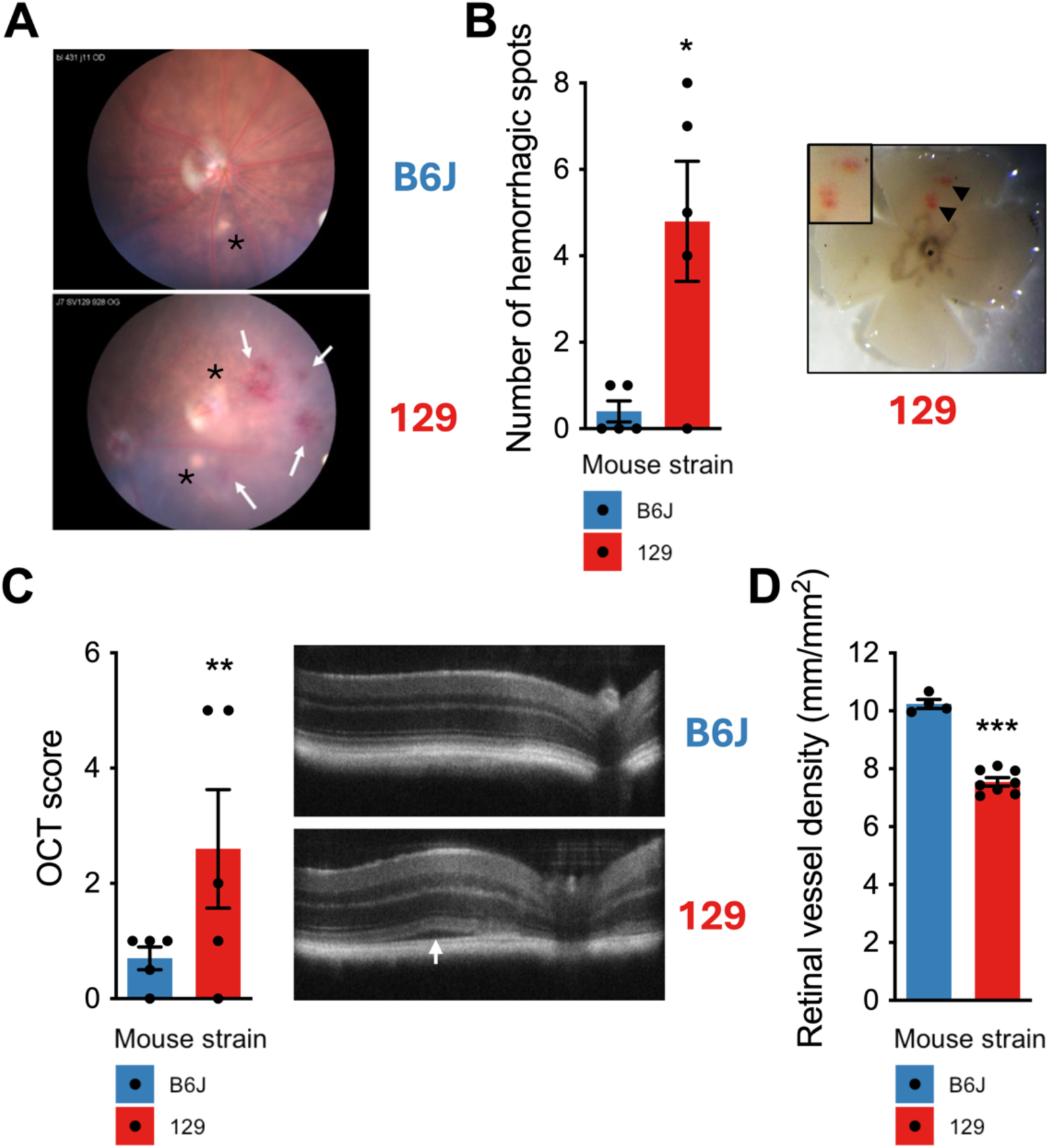
The 129Sv strain develops malignant hypertension with retinal vasculature injury. A- In vivo imaging (microm) performed on 129Sv (129) and C57BL6/J (B6J) mice 8 days after angiotensin-2 pump implantation shows haemorrhage (arrows) and hard exudates (asterisks). Haemorrhagic spots are only found on the 129Sv eye fundus and density of hard exudates is increased in 129 mice compared to B6J mice. B-Haemorrhagic spots (arrow heads) in retinas at day 7 of hypertensive challenge and associated quantification. Values represent mean ± SEM of 5 mice per group. * p < 0.05. C- Subretinal swelling assessed by OCT at day 7 of hypertensive challenge and associated quantification. The white arrow shows retinal detachment. D- Retinal capillary density assessed at day 7 of hypertensive challenge. Representative images of the retina vascular network stained with lectin antibody. Values represent mean ± SEM of 4 B6J mice and 8 129 mice. *** p < 0.001.

Similarly, we observed elevated hematocrit levels at day 7 in hypertensive 129Sv mice, suggestive of hemoconcentration and plasma leakage (Fig. 4A). This was confirmed by elevated wet-to-dry weight kidney weight ratios (Fig. 4B) and by increased vascular permeability to Evans Blue (Fig. 4C).

**Figure 4:**
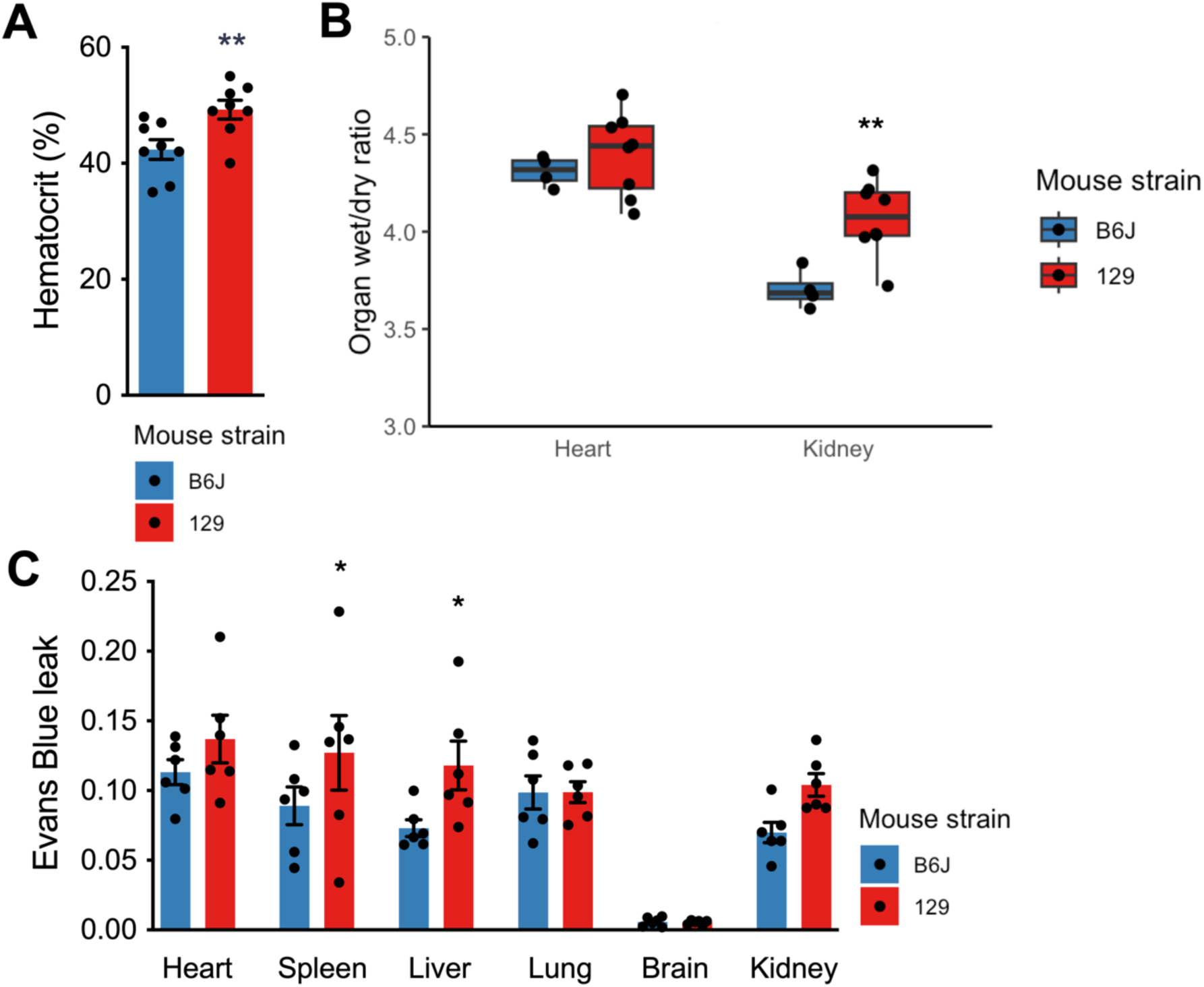
The 129Sv strain develops malignant hypertension with capillary leakage and organ edema. A- Blood hematocrit levels at day 7 of hypertensive challenge in 129Sv (129) mice and C57BL6/J (B6J) mice. Values represent mean ± SEM of 8 mice per group. ** p < 0.01. B- Heart and kidney wet-to-dry weight ratios at day 7 of hypertensive challenge. Values represent mean ± SEM of 4 B6J mice and 8 129 mice. ** p < 0.01. C- Heart, spleen, liver, lung, brain and kidney Evans Blue permeability assay at day 7 of hypertensive challenge. Values represent mean ± SEM of 6 mice per group. * p < 0.05.

### Hypertensive 129Sv mice display altered cardiac and renal hemodynamics and life-threatening arrhythmias

Noninvasive ultrasound assessment of hypertensive mice at day 14 (Fig. 5A) of hypertension onset revealed heart failure in 129Sv mice, as demonstrated by decreased stroke volume and decreased cardiac output. Renal blood flow was accordingly decreased in those mice. Continuous ECG monitoring revealed progressive onset of electrical alterations (Fig. 5B) and arrhythmias (Fig. 5C and Supp. Fig. 2) in hypertensive 129Sv mice during the hypertensive challenge, beginning on day 2. On both days 2 and 12, 129Sv mice exposed to Ang II exhibited a ∼15% prolongation of the QTc from a baseline of 53.7 ± 2.7ms (p < 0.05 *vs.* baseline). In addition, SDNN, as a global variability index, is dramatically reduced 12 days after initiation of Ang II administration, revealing a profound autonomic dysfunction (Fig. 5C). Such a decrease is a strong, independent predictor of mortality because it reflects the heart’s ability to adapt to physiological and pathological stressors and so, is commonly associated with a higher risk of cardiac events and death. At baseline, the frequency of ventricular ectopy (VE) was low (3.8 ± 0.6 events/3h), and mice treated with Ang II demonstrated increased VE (day 2: 10.7 ± 1.5; day 12: 15.4 ± 1.9 events/3h). 129Sv mice exposed to Ang II also displayed electrical alternans at day 2 (100% of mice) and day 12 (80% of mice). Finally, evidence of ventricular tachycardia (VT) and atrial fibrillation (AF) was observed in 60% of mice receiving Ang II on day 12. To note, 2 out of 7 hypertensive 129Sv mice died from cardiac arrest, as illustrated in Supplementary Figure 2. These observations suggest that heart disease may be a cause of death in hypertensive 129Sv mice.

**Figure 5:**
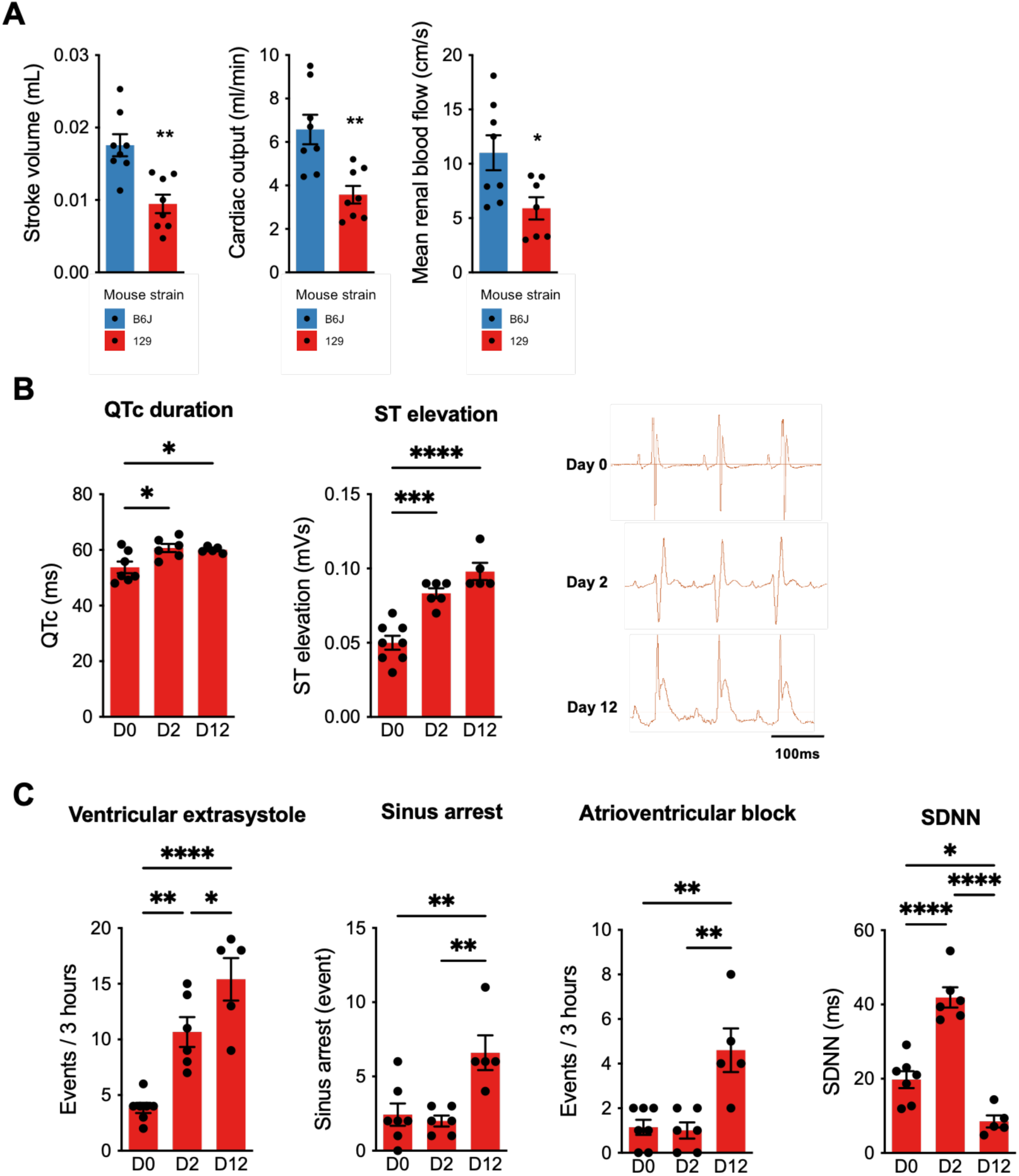
The 129Sv strain develops malignant hypertension with cardiac insufficiency and arrhythmia. A- Noninvasive hemodynamics ultrasound assessment at day 14 of hypertensive challenge in 129Sv (129) mice and C57BL6/J (B6J) mice. Values represent mean ± SEM of 8 mice per group. ** p < 0.01. B- Electrical alterations at day 0, day 2 and day 12 of hypertensive challenge in 129 mice. Left panels: mean values for ST elevation and QTc duration. Right panels: representative ECG traces. Values represent mean ± SEM of 7 mice at day 0, 6 mice at day 2 and 5 mice at day 12 * p < 0.05, ** p < 0.01. C- Frequency of arrhythmic events (number of events per 3 hours) at day 0, day 2 and day 12 of hypertensive challenge in SV129 mice. Values represent mean ± SEM of 7 mice at day 0, 6 mice at day 2 and 5 mice at day 12. * p < 0.05, ** p < 0.01, *** p<0.001, **** p<0.0001

### Multiorgan microvascular disease in hypertensive 129Sv mice is associated with an imbalance in angiogenic factor

The systemic microvascular disorder displayed by hypertensive 129Sv is reminiscent of preeclampsia, a hypertensive disorder of pregnancy with clinical features closely resembling those of HTEM. Preeclampsia has been associated with an impaired angiogenic balance thought to stem from markedly elevated circulating sFlt-1 levels. sFlt-1, the soluble form of the VEGFR1 receptor, acts as a VEGFA scavenger in preeclampsia and is secreted by the placenta in this specific pathological context, which cannot be the case here since only male subjects were examined. We observed markedly elevated plasma sFlt-1 levels in 129Sv males after the onset of hypertension at levels measured in preeclampsia, whereas these levels remained low in B6J males (Fig. 6A).

**Figure 6:**
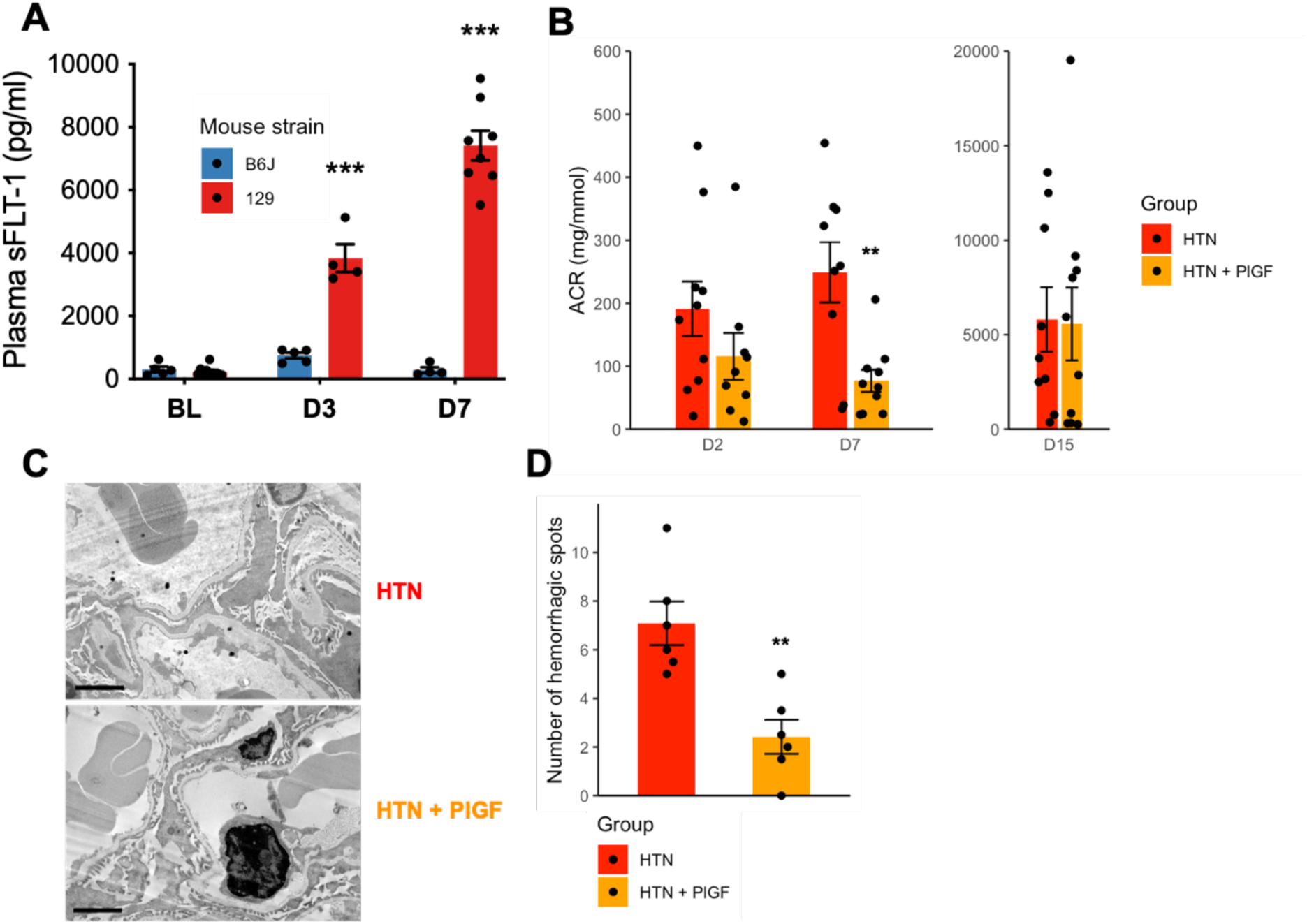
Elevated sFlt-1 levels in 129Sv mice upon hypertensive challenge and PlGF- mediated rescue of renal and retinal phenotypes. A- Plasma sFlt-1 levels in hypertensive B6J and 129S mice. Values represent mean ± SEM of 8 mice per group. ** p < 0.01, *** p < 0.001. B- Urinary albumin-to-creatinine ratios in hypertensive 129S mice with or without rhPlGF-2 supplementation. Values represent mean ± SEM of 10 mice per group. ** p < 0.01. C- Glomerular filtration barrier ultrastructure in electronic transmission microscopy at day 7 of hypertensive challenge. Scale bar = 2 μm. D- Count of retinal haemorrhagic spots in hypertensive 129S mice with or without rhPlGF-2 supplementation. Values represent mean ± SEM of 6 mice per group. ** p < 0.01.

Thus, we hypothesized that elevated circulating sFlt-1 contributed to microvascular impairment by trapping VEGFA, which is necessary for maintening specialized endothelia such as the glomerular endothelium. To test this hypothesis, we delivered subcutaneous rhPlGF-2 to hypertensive 129Sv mice to trap the circulating sFlt-1 and tip the balance toward the pro-angiogenic side. In these rhPlGF-2-supplemented mice, we observed a partial reversal of the hypertensive disease phenotype with decreased albuminuria (Fig. 6B), preserved GFB ultrastructure (Fig. 6C), and decreased retinal hemorrhages (Fig. 6D) at day 7 of hypertension onset, demonstrating that dysregulated sFlt-1/VEGFA balance is responsible for 129Sv mice hypertensive microvascular disease.

*Hematopoietic and non-hematopoietic cells are responsible for 129Sv mice’s sensitivity to hypertension*.

We next performed bone marrow (BM) transplantation to investigate whether the susceptibility of 129Sv mice to hypertension-induced microvascular injury and the origin of sFlt-1 are mediated by the immune system. Bone marrow reconstitution efficiency in recipient mice was assessed by flow cytometry using CD45 allelic markers. As shown in Supp. Fig. 3, BM-transplanted mice demonstrated robust engraftment, with approximately 80–85% of cells expressing the donor marker. Renal function was evaluated on day 7 after the hypertensive challenge by measuring BUN, serum albumin, and ACR levels. B6J mice with B6J BM exhibited preserved renal function, whereas 129Sv mice with 129Sv BM developed renal dysfunction. Mice with mismatched BM (B6J mice with 129Sv BM and 129Sv mice with B6J BM) showed intermediate degrees of renal injury, with 129Sv mice receiving B6J BM tending toward more severe dysfunction than B6J mice with 129Sv BM (Fig. 7A–C). These findings indicate that non- hematopoietic cells of 129Sv origin may be the primary contributors to organ injury in Ang II- challenged mice. Additionally, sFlt-1 levels were elevated in both B6J mice with 129Sv BM and 129Sv mice with B6J BM compared to B6J mice with B6J BM, though to a lesser extent than in 129Sv mice with 129Sv BM. This suggests that both hematopoietic and non-hematopoietic 129Sv cells are required to drive sFlt-1 expression (Fig. 7D).

**Figure 7:**
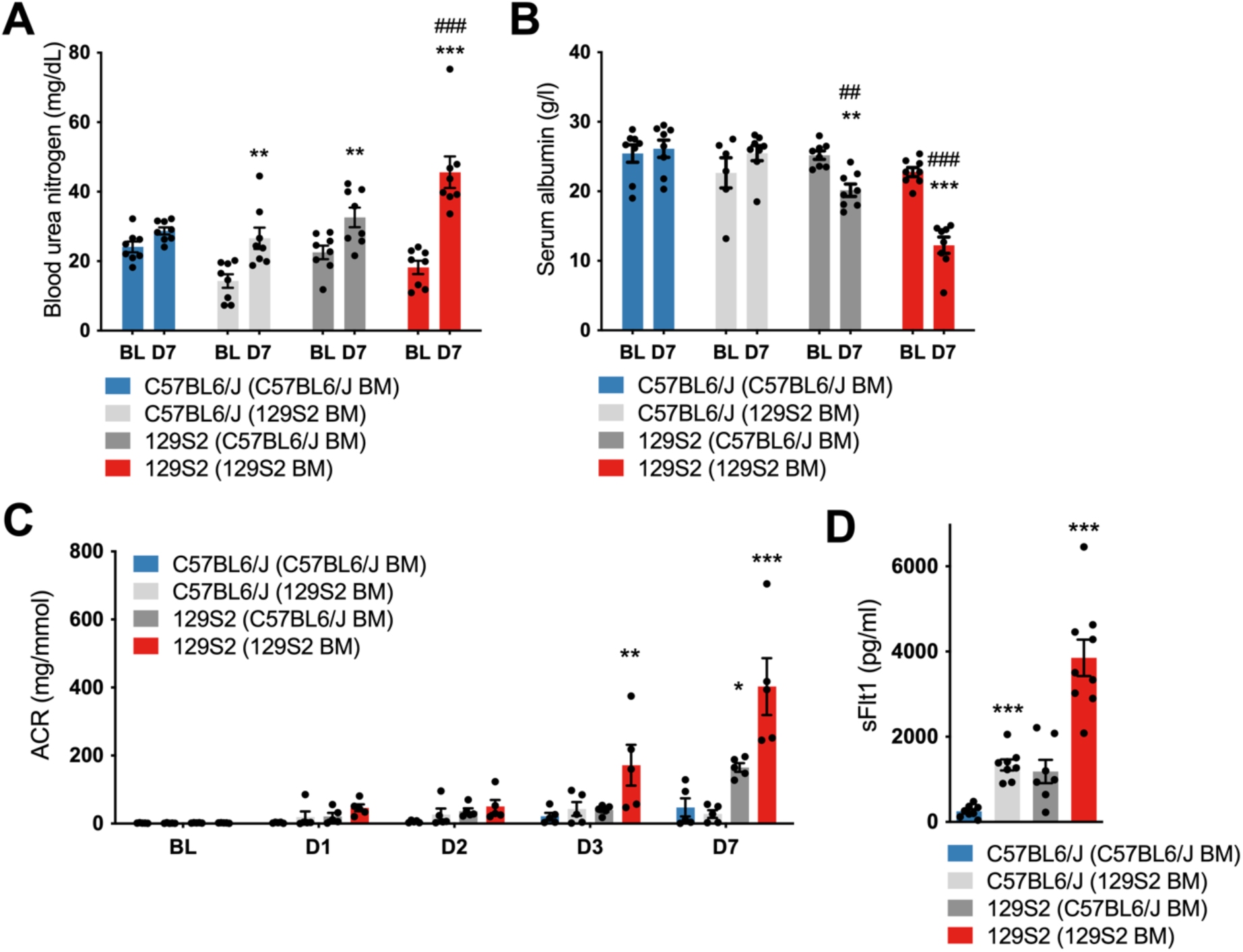
Loss of tolerance to hypertensive stress is mediated both by the immune system and tissue in 129Sv mice. A- Blood urea nitrogen measured at day 7 of hypertensive challenge in C57BL6/J or 129Sv mice recipient of C57BL6/J or 129Sv bone marrow 6 weeks earlier. N= 8 mice per group. Values represent mean ± SEM. 2 way-ANOVA with Tukey post hoc test, ** p < 0.01, *** ### p < 0.001. * for comparison between D7 and respective D0, # for comparison between group to C57BL6/J (C57BL6/J BM). B- Plasma albumin measured at day 7 of hypertensive challenge in C57BL6/J or 129Sv mice recipient of C57BL6/J or 129Sv bone marrow 6 weeks earlier. N= 8 mice per group. Values represent mean ± SEM. 2 way-ANOVA with Tukey post hoc test, ** ## p < 0.01, *** ### p < 0.001. * for comparison between D7 and respective D0, # for comparison between group and C57BL6/J (C57BL6/J BM). C- Urine albumin-to- creatinine ratio across a 7-day hypertensive challenge. N=5 mice per group from baseline (BL) to day (D) 7. Values represent mean ± SEM. 2 way-ANOVA with Tukey post hoc test, * p < 0.05, ** p < 0.01, *** p < 0.001. * represent statistical significance between a group and C57BL6/J (C57BL6/J BM) at a given time. D- Plasma sFlt-1 levels measured at day 7 of hypertensive challenge in C57BL6/J or 129Sv mice recipient of C57BL6/J or 129Sv bone marrow 6 weeks earlier. Values represent mean ± SEM of 8 mice per group. Welch ANOVA with Dunnett’s T3 post hoc test, *** p < 0.001.

### Hypertension impairs glomerular cell function with major impacts on glomerular endothelial cells

We performed single-cell RNA sequencing of glomeruli from 129Sv mice, either untreated (129) or after a 7-day hypertensive challenge with or without PlGF treatment (129_Ang2 and 129_Ang2_PlGF) (Fig. 8A). Following segmentation of glomerular endothelial cells (GEnCs), podocytes, and mesangial cells (Fig. 8B and Supp. Fig. 4A-B), we conducted differential gene expression (DEG) analysis between conditions and performed pathway enrichment analysis (Fig. 8C-D). Our results show that Ang II, combined with a high-salt diet, predominantly affects the transcriptome of GEnCs. The main impact involves a downregulation of transcription and translation processes in GEnCs. This leads to profound alterations in essential endothelial functions, including angiogenesis, coagulation, and glomerular filtration. Pathways including VEGFR signaling, hypoxia response, cell death, and autophagy are modulated, highlighting the significant impact of hypertension-induced stress on cell survival. Interestingly, we observed a decrease in the endothelial inflammatory signature, particularly involving the NF-κB pathway, interferon signaling, and the IL-1 response (Fig. 8E-G and Supp. Fig. 4C-D).

**Figure 8:**
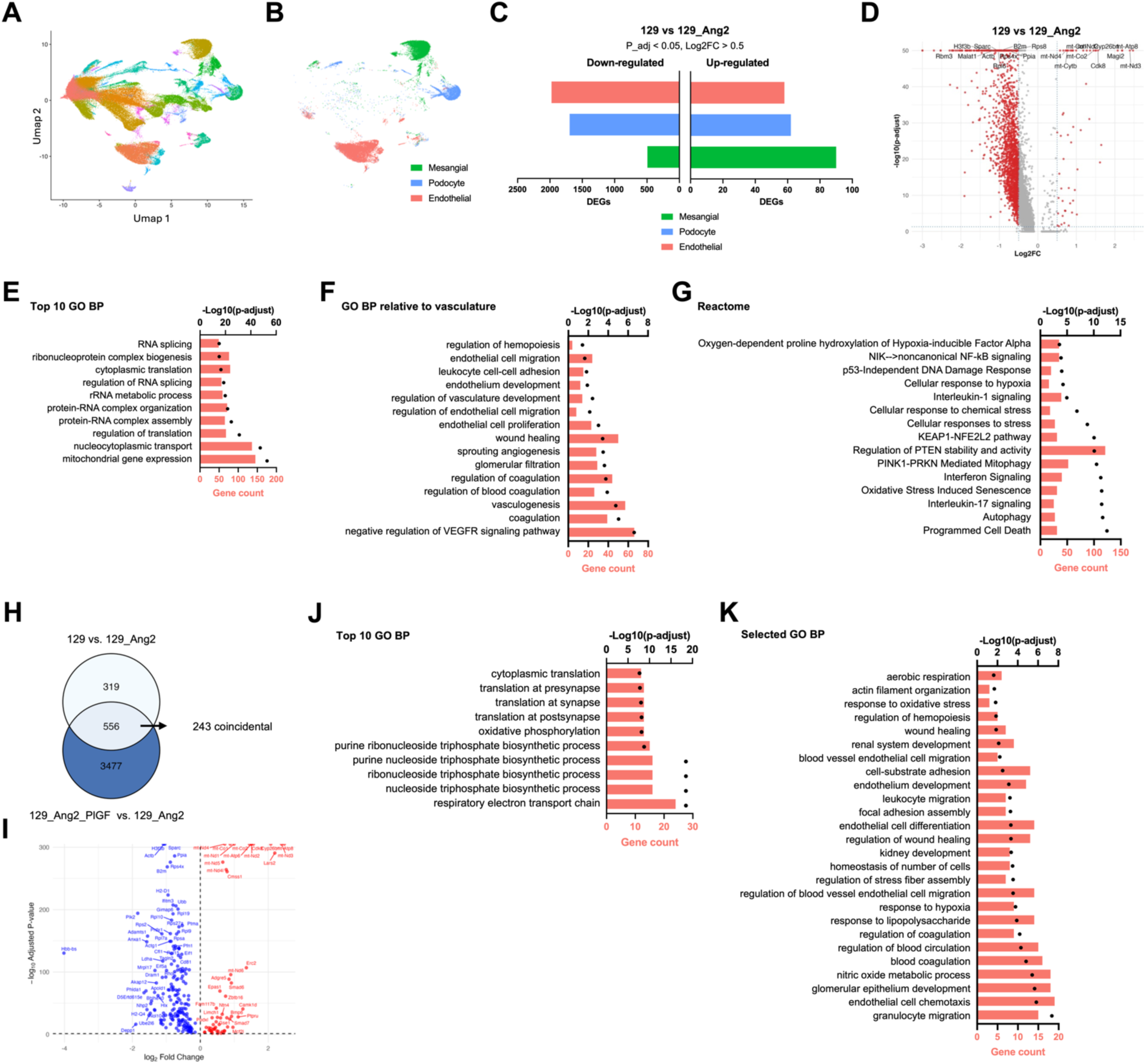
Single cell RNAseq of glomerular cells highlight injury mechanisms in endothelial cells of 129sv mice after hypertensive challenge. A. UMAP plot showing cell clustering. B. UMAP plot showing Endothelial, Podocyte and Mesangial cell clustering. C. Bar graph representing the number of upregulated and downregulated genes in the 129 vs 129_Ang2 comparison. The x-axis shows gene count. D. Volcano plot of differential gene expression between 129 and 129_Ang2. E-G. Bar chart of the top 10 Gene Ontology Biological Process (GO BP) (E), GO BP relative to vasculature terms (F), and selected Reactome (G) terms for differentially expressed genes, with -log₁₀(p-adjusted) values and gene count. H. Venn diagram showing overlap of differentially expressed genes between 129 vs 129_Ang2 and 129_Ang2_PlGF vs 129_Ang2. Numbers indicate unique, shared genes and shared genes with coincidental variation. I. Volcano plot of differential gene expression shared between 129 vs 129_Ang2 and 129_Ang2_PlGF vs 129_Ang2 with coincidental variation. J-K. Top 10 GO BP (J) and selected GO BP (K) terms for differentially expressed genes shared between 129 vs 129_Ang2 and 129_Ang2_PlGF vs 129_Ang2 with coincidental variation, with -log₁₀(p- adjusted) values and gene count.

In podocytes, cellular metabolism appears to be most affected by oxidative stress and hypoxia. Cytoskeletal remodeling signals were also detected, consistent with the effacement of foot processes (Supp. Fig. 5). In mesangial cells, whose transcriptome is less affected than that of podocytes and GEnCs, we also observed reduced translation, oxidative stress, and hypoxia response signals, as well as metabolic alterations. These changes manifest as modifications in ECM organization, cell motility, and TGFβ signaling. Interestingly, pathways such as coagulation and endothelial cell activation were also identified in mesangial cells, underscoring the interactions and cross-communication between glomerular cell types (Supp. Fig. 6).

Finally, we examined the impact of PlGF treatment on glomerular cells. Pathway analysis of DEGs among 129, 129_Ang2, and 129_Ang2_PlGF groups revealed consistent positive or negative fold-changes in gene expression (Fig. 8H-I, Supp. Fig. 5E-F, Supp. Fig. 6E-F). PlGF treatment partially restored essential cellular functions in GEnCs by targeting multiple mechanisms, including protein translation and cellular metabolism (Fig. 8J-K). Notably, podocytes also exhibited improvements in metabolic activity and cytoskeletal organization (Supp. Fig. 5G-H), while mesangial cells showed changes in cellular metabolism, differentiation, growth, and ECM (Supp. Fig. 6G-H). Together, our results demonstrate that PlGF partially prevents hypertension-mediated glomerular injury in 129Sv mice.

## Discussion

In this study, we demonstrate a stark difference in tolerance to severe hypertension between two commonly used mouse strains. While B6J mice exhibit no evidence of acute TOD after severe BP elevation, 129Sv mice with similar telemetry-verified BP develop a lethal syndrome characterized by acute damage to retinas, heart, and kidneys. Thus, severely hypertensive 129Sv mice are a valuable model for studying human HTEM. To the best of our knowledge, this novel mouse model is the first to recapitulate the human condition closely. In particular, this is the first mouse model of hypertensive retinal hemorrhage and sudden death to our knowledge. While an HTEM phenotype has previously been noted in various rat models of hypertension, such as nitro-L-arginine-methylester-treated rats or specific genetic strains (14), mouse models of hypertension typically do not follow a malignant course and more closely model the consequences of chronic BP elevation in humans (30, 31). The wealth of genetic tools readily available for studying mouse models (32) makes this novel HTEM mouse model a valuable asset for experimentally dissecting the mechanisms of the human disease.

While genome-wide association studies in humans have identified multiple genetic polymorphisms associated with BP levels (33–35), little is known about the genetic determinants of TOD. Our comparison of B6J and 129Sv strains’ tolerance to severe hypertension unequivocally demonstrates that BP-independent genetic factors govern the development of TOD in this context. Likewise, a recent report described increased cardiovascular and kidney damage after 5/6 nephrectomy in 129Sv mice (36). Our HTEM model may therefore serve as a useful platform for the mechanistic study of susceptibility to hypertension-induced TOD. Compared with the B6J strain, the 129Sv strain has a two-renin gene background, is salt-sensitive (37), and has decreased tolerance to the deoxycorticosterone acetate/salt model of hypertension (21). 129Sv mice also exhibit an impaired aortic response to acetylcholine (38) and display markedly aggravated glomerular injury with glomerular thrombotic microangiopathy (TMA) lesions in the nephrotoxic serum- induced crescentic glomerulonephritis model (39) as compared to C57Bl6/J. In the latter case, glomerular expression profiling, using microarrays and Western blot analysis in B6J TMA- resistant and 129Sv TMA-prone mice, demonstrated major differences in vascular endothelial growth factor (VEGF)/VEGF receptor (VEGFR) 2 pathways, despite similar *Vegfa* expression levels. How these specific differences contribute to the development of HTEM in our Ang II and high-salt diet-induced model remains to be studied.

Strikingly, we observed markedly elevated levels of circulating sFlt-1 in 129Sv mice challenged with severe hypertension. sFlt-1, the circulating splice variant of VEGFR1, binds free VEGFA and acts as an anti-angiogenic factor as it sequesters this ligand and thus prevents the binding to and subsequent signal transduction of receptors with pro-angiogenic signaling properties, such as VEGFR2. VEGFA-VEGFR2 signaling is essential for maintaining endothelial integrity, vasodilation and permeability (40), especially in specialized fenestrated endothelia such as the glomerular endothelium (41, 42). Preeclampsia, a hypertensive disorder of pregnancy with clinical features closely resembling those of HTEM, is associated with increased plasma sFlt-1 stemming from placental sFlt-1 overexpression, and decreased plasma PlGF, a proangiogenic ligand selectively binding to sFlt-1 (43, 44). The pathogenetic role of this dysregulated angiogenic balance on disease manifestations was further demonstrated in various animal models of preeclampsia (45, 46). Indeed, adenoviral delivery or transgenic overexpression of human or murine soluble Flt1 (and, in some models, soluble Endoglin) in pregnant mice induces dose-dependent gestational hypertension, proteinuria, glomerular endotheliosis, podocyte injury, and endothelial dysfunction (43, 44, 47). Interestingly, adenoviral delivery of high sFlt1 concentrations does not induce HTEM in non-pregnant, non-hypertensive C57Bl6/J females and males, suggesting that additional cofactors are required to reveal microvascular sensitivity to impaired VEGFR signaling. In our work, only males were studied, illustrating the possibility of extraplacental sFlt-1 overexpression during severe hypertension. Bone marrow transplantations suggested that both hematopoietic and nonhematopoietic cells may contribute to the observed plasma elevation. Indeed, VEGFR1 expression has been demonstrated in several hematopoietic cell types (48–50), including monocytes in ANCA vasculitis (51), raising the question of an interaction between the immune system and the VEGF-sFlt-1 balance. The bone marrow transplant experiments indicated that increased sFlt-1 levels in HTEM mice may originate both from the bone-marrow-derived cells and the tissues, or that a reciprocal interaction between immune cells and the vasculature in a prone HTEM background is required for full-blown disease.

Indeed, elevated levels of circulating sFlt-1 may stem from other tissues, such as adipose tissue (52), and, above all, from endothelial overexpression or shedding (53). In this context, it is debatable whether sFlt-1 elevation in our model reflects a pathogenetic mechanism or is merely a bystander phenomenon secondary to endothelial damage. When treating hypertensive 129Sv mice with rhPlGF-2 to restore pro-angiogenic signaling, we observed a significant partial reversal of the disease phenotype and glomerular endothelial cells transcriptomic alterations, providing evidence that excess sFlt-1 is a causal contributor to microvascular damage in this HTEM model. These results highlight potential mechanistic links between preeclampsia and HTEM and should prompt increased scrutiny of the angiogenesis balance during HTEM. The only study focusing on sFlt-1 levels in HTEM patients demonstrated only a slight increase as compared to patients with nonmalignant hypertension (54). This elevation was far below that observed in preeclampsia and persisted at the 12-month follow- up. Further research is therefore needed to assess whether excess sFlt-1 may be a causal contributor to TOD during HTEM, or to identify alternative or additive mechanisms of dysregulated angiogenesis in this specific context.

In conclusion, we describe a novel Ang II and high-salt diet-induced model of HTEM in mice, highly reminiscent of human disease, which we hope will facilitate future investigations into the mechanisms of this poorly understood condition. Interestingly, we uncovered a dysregulated balance of circulating angiogenic factors in 129Sv mice suffering from HTEM. This finding suggests pathophysiological similarities between human preeclampsia and HTEM and warrants further investigation of pro- and anti-angiogenic factors during HTEM.

## Perspectives

The present work provides the first description of a mouse model that faithfully recapitulates the acute, multi-organ manifestations of hypertensive emergency. This model opens new avenues for research into the mechanisms driving target organ damage during abrupt blood pressure elevation. Our findings highlight the contribution of BP-independent, genetically encoded determinants of vascular fragility, pointing to the need for renewed focus on inter-individual susceptibility rather than blood pressure thresholds alone. The discovery of a dysregulated VEGFA/sFlt-1 axis as a causal contributor to endothelial injury suggests that angiogenic imbalance may represent a broader paradigm in severe hypertensive syndromes. Future work should identify the cellular sources and upstream triggers of sFlt-1 overproduction during hypertensive stress, and determine whether modulating angiogenic pathways can prevent or reverse microvascular injury.

## Novelty and Relevance

### What Is New?

- Identification of the first mouse model that reproduces the multi-organ microvascular injury characteristic of hypertensive emergency.
- Discovery that hypertensive injury severity depends on genetic background despite similar blood pressure levels.
- Demonstration that angiogenic imbalance drives endothelial dysfunction and target organ damage.

### What Is Relevant?

- Highlights that BP-independent mechanisms determine hypertensive emergency susceptibility.
- Reveals angiogenic imbalance as a key pathogenic pathway in severe hypertensive states.
- Provides a relevant model for mechanistic and therapeutic studies.

### Clinical/Pathophysiological Implications?

- This work suggests that susceptibility to hypertensive emergency is genetically determined and involves dysregulated VEGFA/sFlt-1 signaling.
- Targeting angiogenic imbalance may offer new therapeutic opportunities to prevent acute target organ damage in severe hypertension.

## Supplementary Figures

**Supplementary Figure 1:**
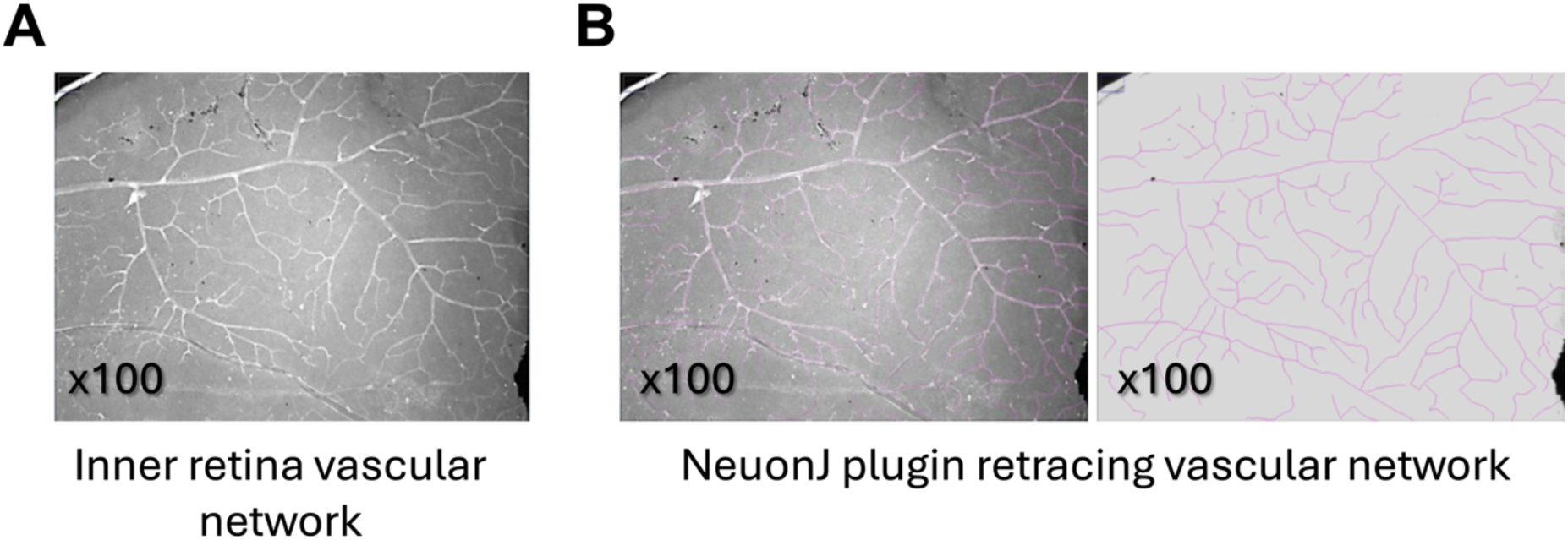
Retinal vascular density. Representative images of retina vascular network stained with lectin antibody (A). Vascular network density was obtained using NeuronJ plugin on ImageJ software (B).

**Supplementary Figure 2:**
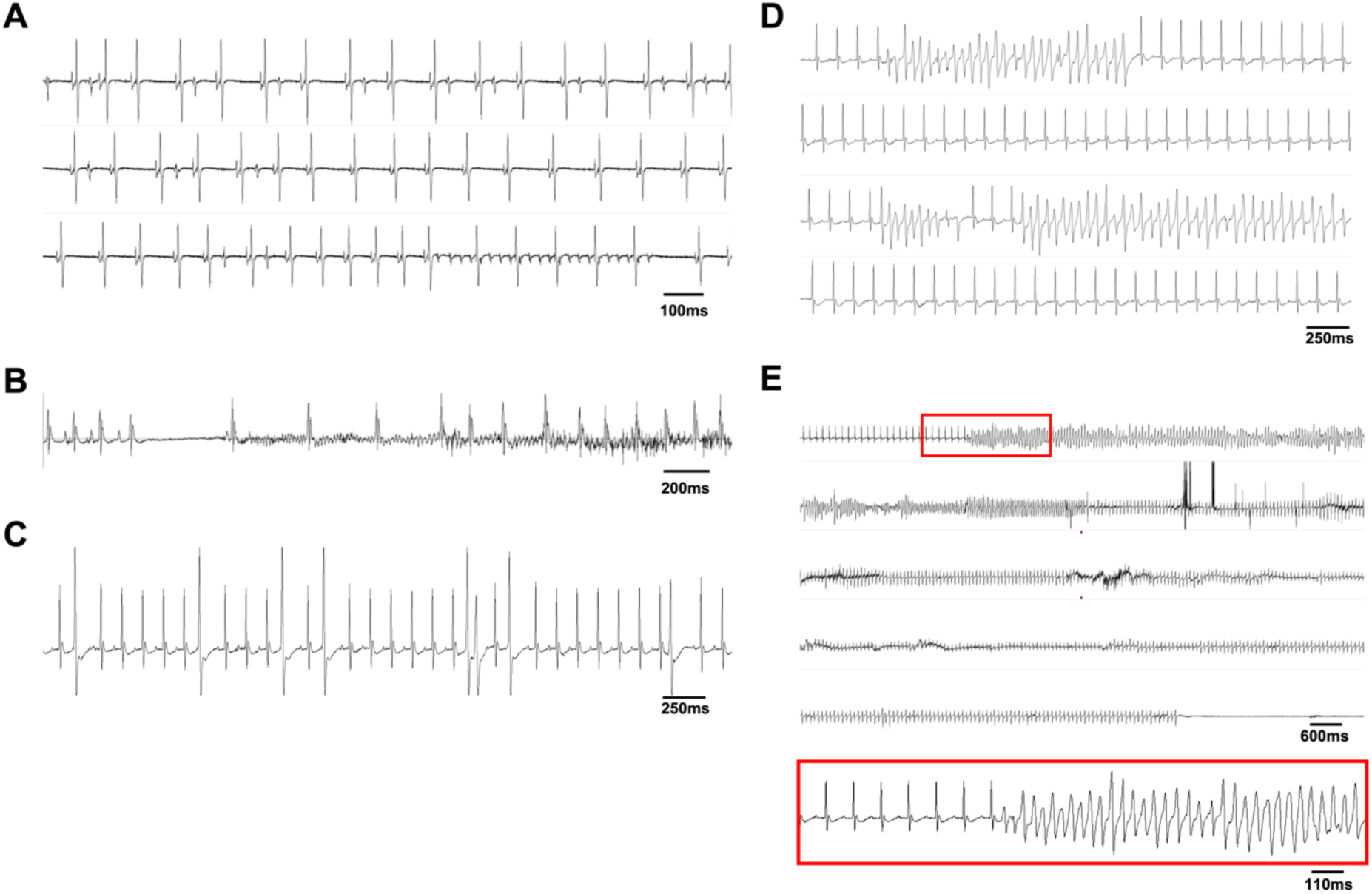
Lethal cardiac arrhythmia in 129Sv mice upon hypertensive challenge. Typical ECG changes seen in SV129 mice treated for 2 weeks with angiotensin II (n = 5-7 mice/group. A. episode of atrial flutter B. atrial fibrillation. C. ventricular ectopy seen in the Ang II group; D. short run of ventricular tachycardia; E. sustained ventricular tachycardia degenerating in ventricular fibrillation and cardiac death.

**Supplementary Figure 3:**
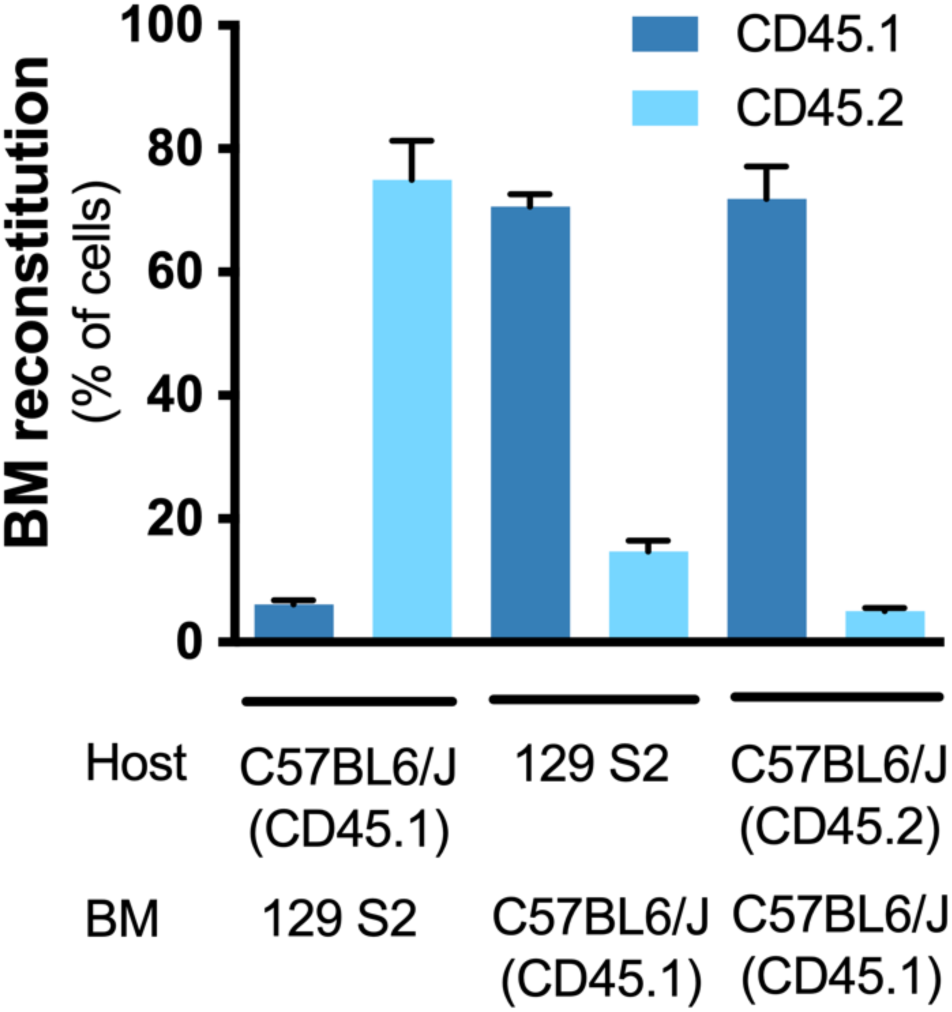
Bone marrow transplantation efficacy. Bone marrow reconstitution efficiency was assessed by flow cytometry.

**Supplementary Figure 4:**
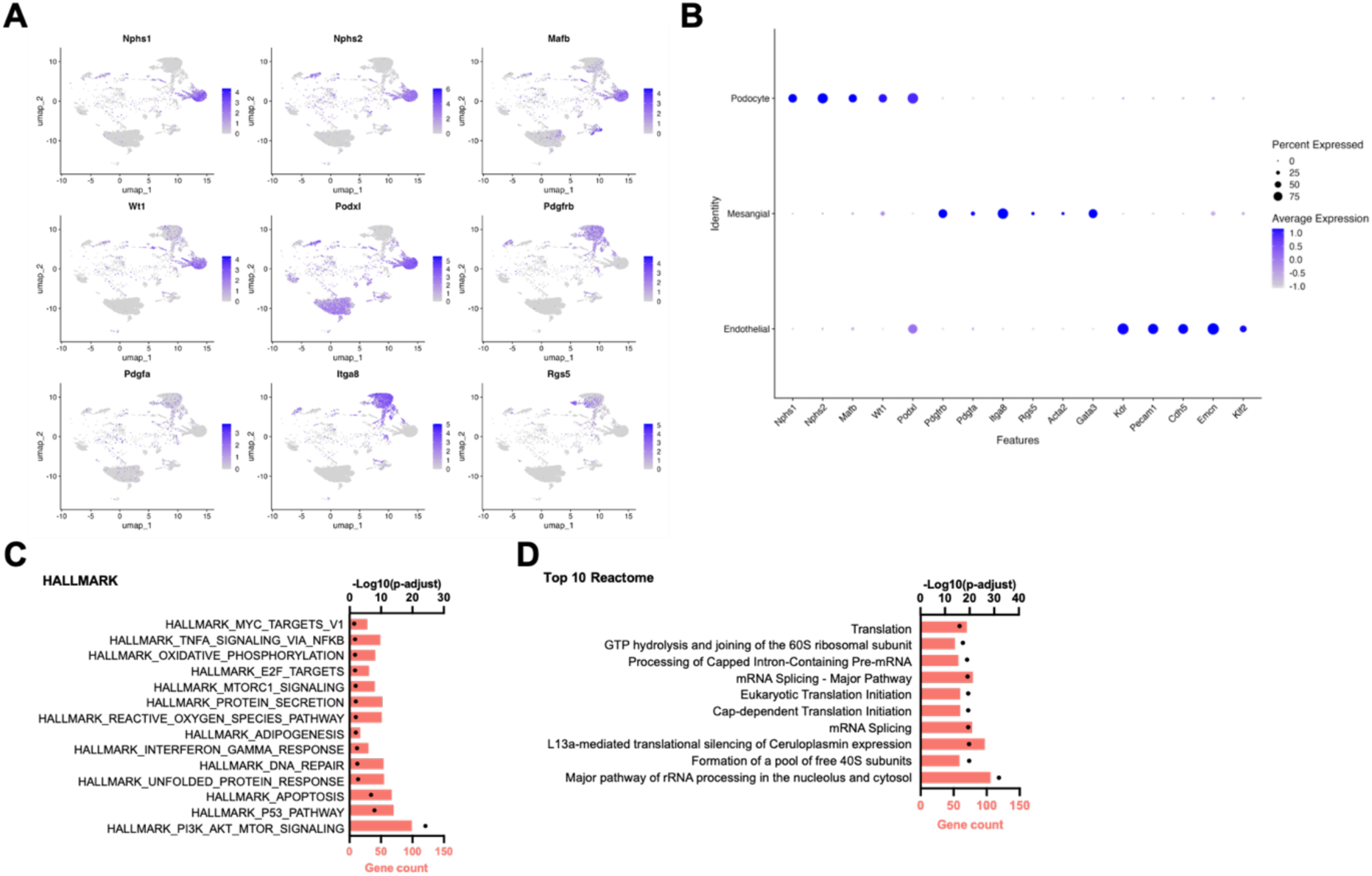
Extended. Figure 7**, single-cell RNA-seq of glomerular cells highlights injury mechanisms in endothelial cells of 129Sv mice after hypertensive challenge.** A. Scatter plots showing the expression levels of reference genes across cells. The color gradient from gray to purple indicates increasing gene expression. B. Dot plot representing the percentage of cells expressing each gene across different cell types. Dot size reflects the proportion of expressing cells, while color intensity indicates average expression levels. C-D. Bar graph showing the top enriched hallmark gene sets (C) and Top 10 Reactome pathways (D) for differentially expressed genes between 129 and 129_Ang2.

**Supplementary Figure 5:**
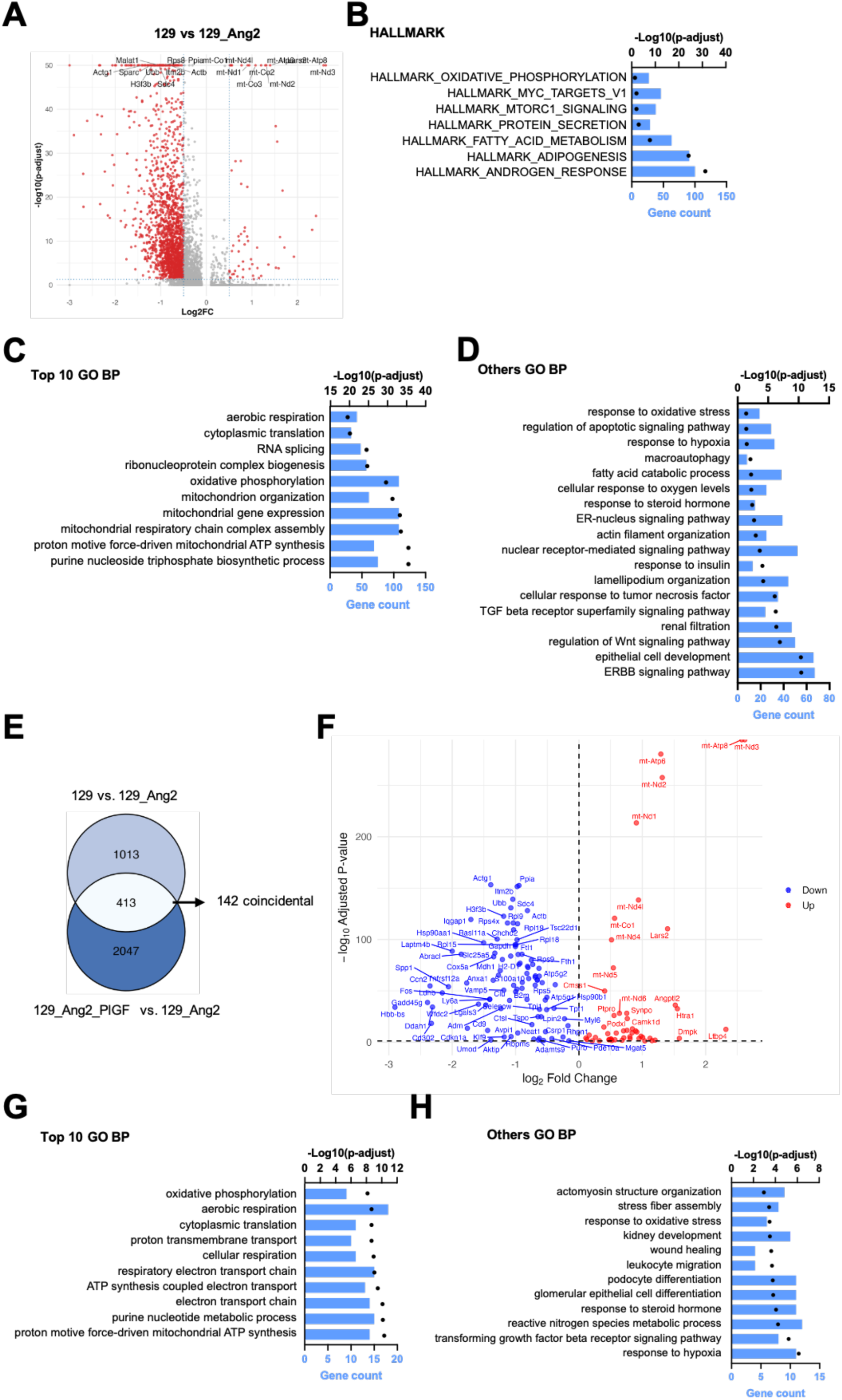
Single cell RNAseq of glomerular cells highlight injury mechanisms in podocytes of 129sv mice after hypertensive challenge. A. Volcano plot of differential gene expression between 129 and 129_Ang2. B-D. Bar chart of the HALLMARK (B), top 10 Gene Ontology Biological Process (GO BP) (C), and selected GO BP (D) terms for differentially expressed genes, with -log₁₀(p-adjusted) values and gene count. E. Venn diagram showing overlap of differentially expressed genes between 129 vs 129_Ang2 and 129_Ang2_PlGF vs 129_Ang2. Numbers indicate unique, shared genes and shared genes with coincidental variation. F. Volcano plot of differential gene expression shared between 129 vs 129_Ang2 and 129_Ang2_PlGF vs 129_Ang2 with coincidental variation. G-H. Top 10 GO BP (G) and selected GO BP (H) terms for differentially expressed genes shared between 129 vs 129_Ang2 and 129_Ang2_PlGF vs 129_Ang2 with coincidental variation, with -log₁₀(p-adjusted) values and gene count.

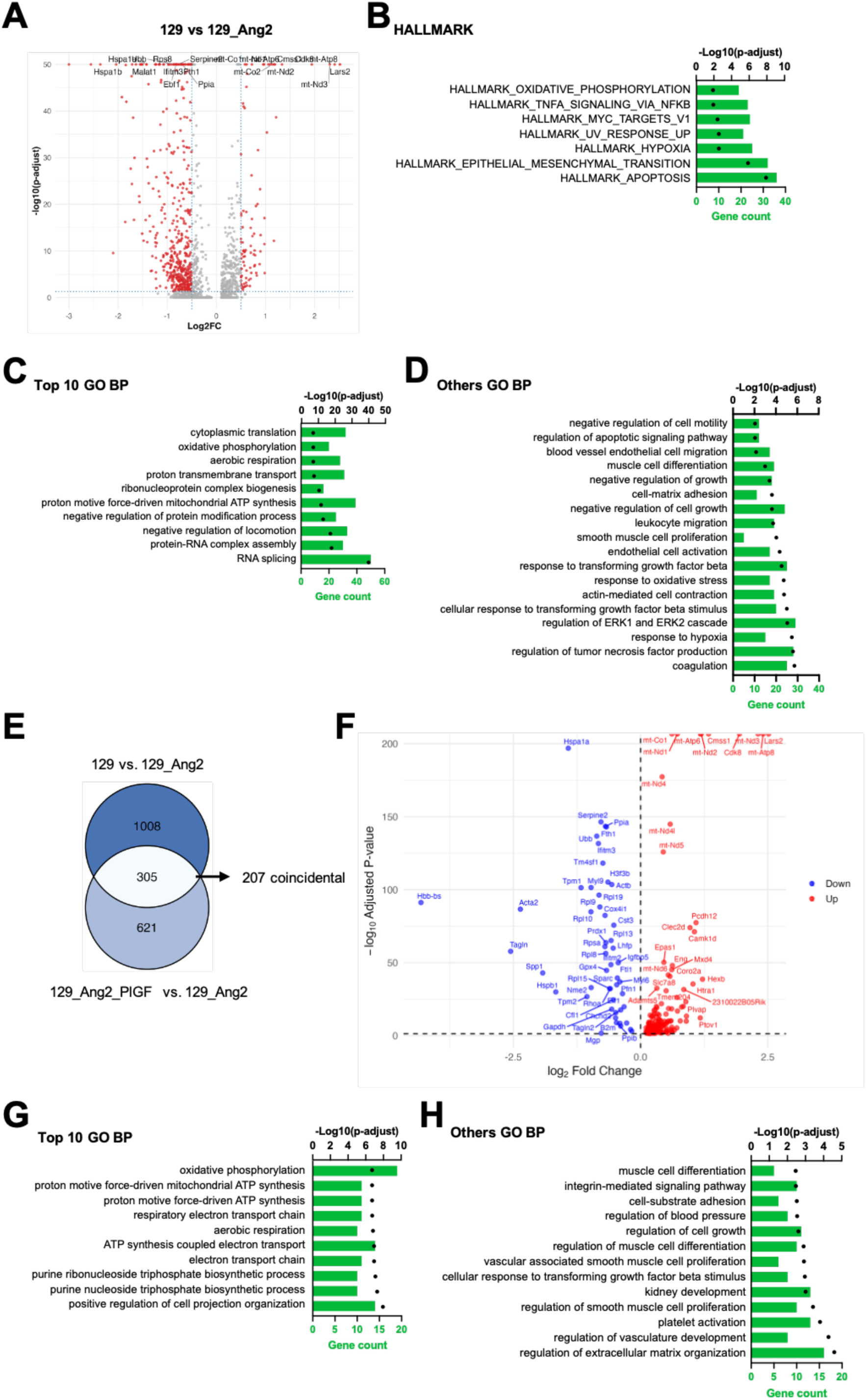

## Acknowledgments

We thank Corina Suldac, Gwendoline Luque and the ERI 970 team for assistance in animal care and handling. We thank Véronique Oberweis and Philippe Coudol for administrative support. We also thank Alain Schmitt from Electronic microscopy platform at the Cochin Institute. We thank Nicolas Sorhaindo from the biochemistry platform at Bichat hospital. We thank Brigitte Izac and Sébastien Jacques from the Cochin Institute Genomic platform.

## Data availability

All data supporting the findings of this study are included in the article and its supplementary information files. Single-cell RNA-sequencing datasets generated during this study will be deposited in a public repository and made accessible at the time of publication. Any additional information or materials related to this work are available from the corresponding authors upon reasonable request.

## Abbreviations

ACR: albumin-to-creatinine ratio
AF: atrial fibrillation
AngII: angiotensin II
ANOVA: analysis of variance
BM: bone marrow
BP: blood pressure
BSA: bovine serum albumin
BUN: blood urea nitrogen
B6J: C57BL/6J mouse
CBC: complete blood count
CD45: cluster of differentiation 45
DEG: differentially expressed genes
ECG: electrocardiogram
ECM: extracellular matrix
EDTA: ethylenediaminetetraacetic acid
FCS: fetal calf serum
GEM: gel bead-in emulsions
GEnCs: glomerular endothelial cells
GFB: glomerular filtration barrier
HBSS: Hank’s balanced salt solution
HTEM: hypertensive emergency
i.v.: intravenous
PBS: phosphate-buffered saline
rhPlGF-2: recombinant human PlGF-2
SD: standard deviation
SEM: standard error of the mean
sFlt-1: soluble fms-like tyrosine kinase-1
STE: ST elevation
TOD: target organ damage
UMAP: uniform manifold approximation and projection
VE: ventricular ectopy
VEGFA: vascular endothelial growth factor A
VEGFR: vascular endothelial growth factor receptor
VT: ventricular tachycardia
129sv: 129S/SvPasCrl mouse

